# Joint consideration of selection and microbial generation count provides unique insights into evolutionary and ecological dynamics of holobionts

**DOI:** 10.1101/2024.11.13.623479

**Authors:** William S. Pearman, Allen G. Rodrigo, Anna W. Santure

## Abstract

The relationship between, and joint selection on, a host and its microbes – the holobiont – can impact evolutionary and ecological outcomes of the host and its microbial community. Here we present a novel agent-based modelling framework for understanding the ecological dynamics of hosts and their microbiomes. Our model explicitly incorporates numerous microbial generations per host generation allowing for selection on both host and microbe. We apply our model to explore community fitness and diversity in the face of rapid environmental change. We demonstrate that multiple microbial generations can buffer changes experienced across host lifetimes by smoothing environmental transitions. Our simulations reveal that microbial fitness and host fitness may be at odds with each other when considering the impact of vertical inheritance of microbial communities from a host to its offspring – where high values favour microbial fitness, while low values favour host fitness – these tradeoffs are minimized when microbial generation count per host generation is high. We suggest that these results arise from ‘cross-generational priority effects’ which maintain diversity within the community and can subsequently be acted upon by selection. Our model is readily extensible into new areas of holobiont research and provides novel insights into holobiont evolution under variable environmental conditions.

## Introduction

Over recent years there has been an increasing interest in understanding host-microbiome co-evolution and the potential for the formation of a ‘holobiont’ (Bruijning et al., 2022; Henry et al., 2021; Roughgarden, 2023; Zeng et al., 2015). This interest has been largely prompted by the increasing awareness of the role of microbiomes in influencing host biology, and the idea that microbes, as a result of their shorter generation times, may adapt faster to changing environments and in turn facilitate host adaptation to these changing environments (Petersen et al., 2023; Shah et al., 2024). Although the term ‘holobiont’ is somewhat loaded and complex, with a short but controversial history (Moran & Sloan, 2015), it is generally used to describe a host and its microbiome as a joint unit upon which natural selection can act (Rodrigo, 2023). Because the microbiome can play a role in shaping host fitness (Henry et al., 2021), there is the potential for multilevel selection wherein natural selection operates on both the microbiome, the host, and the joint ‘unit’ (van Vliet & Doebeli, 2019). This idea of multi-level selection with regards to the holobiont has been well modelled through a range of different approaches, with notable advances made by Zeng et al., (2015, 2017), van Vliet and Doebeli (2019), Roughgarden, (2022), and Bruijning et al., (2022). These works largely build off each other and serve to provide a strong theoretical framework for understanding holobiont evolution and ecology.

Holobiont theory is often centered around the mode of transmission of microbes, or where a host obtains their microbiome from. This is understandably of notable importance as traditional or neo-Darwinian evolutionary theory anticipates a lineal inheritance pattern for the evolution of traits. This ‘lineal’ inheritance is referred to as vertical transmission, and contrasts with horizontal transmission, which is the acquisition of microbes from other sources (most commonly the environment). However ‘mixed mode transmission’, where hosts acquire some proportion of their microbiome via vertical transmission, and the remainder from the environment, is likely the dominant form of microbial acquisition (Ebert, 2013).

In this paper, we consider a holobiont as an ecological unit which participates in the evolutionary process, but is not *sensu stricto* a unit of selection (see (Suárez & Stencel, 2020; Veigl et al., 2022) for comprehensive discussions of the holobionts participation in the evolutionary process). In our case we define a holobiont as the relationship between a macroorganism and a community of microbes whereby both host and microbe fitness are impacted by their relationship, and selection can operate on host, microbe, and their relationship. We define holobiont in this sense following the argument from Roughgarden (2020 & 2022), who suggests that 1) holobionts can arise in the absence of strict vertical transmission, and that 2) “ holobiont formation is a unique and simultaneous combination of both evolutionary and ecological processes” (Roughgarden, 2023). Finally, although previous agent based model approaches do not necessarily refer to their systems as holobionts (Bruijning et al., 2022; Zeng et al., 2017), there is an implicit argument that this what they are modelling (Rodrigo, 2023). Thus, we prefer to be explicit in what we model and how we define a holobiont.

As noted by Rodrigo (2023), there are three broad types of models of holobiont evolution. The first, by van Vliet & Doebeli (2019) makes use of multilevel selection theory to understand holobiont evolution and is expressed as a set of mathematical equations rather than as an agent based process. van Vliet & Doebeli (2019) suggest that microbes can evolve altruistic traits under short host-generation times and high levels of vertical inheritance – even when there is a cost to the microbe. Counter to this, Roughgarden (2020) and (2022) propose that a ‘collective’ inheritance of microbes from the environment can lead to the formation of a holobiont through host filtering/selection for environmental microbes, suggesting that vertical transmission is not essential for holobiont formation. Finally, the last set of models are agent based processes first pioneered by Zeng et al., (2015, 2017, 2018) with commonalities with those models of Bruijning et al., (2022) and Daybog & Kolodny (2023). Zeng et al (2015, 2017) developed a model which includes both host and microbial selection, this work showed that host selection alone is not sufficient to suppress microbial diversity while microbial selection can suppress diversity alone. The model developed by Zeng has been subsequently extended by Bruijning et al (2022) and Daybog & Kolodny (2023). Bruijning et al (2022) demonstrate that the fitness benefits to a host from vertical transmission are dependent on both the predictability and variance of an environment. Daybog & Kolodny (2023) demonstrated in their version of the model that microbial beta diversity can remain high, despite the intuition that communities should converge on the ‘optimal’ composition, as a result of a range of factors such as population size and microbial community richness. These agent-based models are sufficiently flexible to incorporate an arbitrary number of hosts and microbes and are readily extendable into new areas of holobiont research – therefore the model we present here is based on an extension of these models.

Our work has been prompted by the observation that there are two core assumptions regularly made with regards to holobiont simulations, the first is that of neutrality, where microbial communities are neutrally assembled within the host without consideration of selection within the host. The second assumption is that there are one or few microbial generations per host generation. Indeed, the assumption regarding singular microbial generations per host generations, despite being practically useful for simulations, ignores an oft-cited argument for the benefits of a microbiome – that multiple microbial generations per host generation can facilitate adaptation of the host in changing environments (Ferreiro et al., 2018; Kolodny & Schulenburg, 2020). Bruijning et al, (2022) simulate up to eight microbial generations within a host generation – yet assume that microbial communities are neutrally assembled within the host, thus for selection to operate on the microbiome it must be indirect through host reproduction. Making such assumptions can be advantageous, both with regards to simplicity but also with regards to computational feasibility (e.g. (Munoz et al., 2018)). However, jointly considering the role of neutrality and multiple microbial generations is an important step towards understanding holobiont evolution (Sieber et al., 2019), particularly because the fitness of both hosts and microbes must benefit from the relationship (or at least not be negatively affected) in order for a holobiont to evolve.

Another simplification of many current models of holobiont evolution is that most models to date have been based on hosts acquiring microbes from a combination of a ‘shedding’ contribution from hosts into the environment and a fixed environmental pool of microbes (Roughgarden, 2023; Zeng & Rodrigo, 2018). In contrast, environments are likely to, in addition, have a substantial proportion of resident microbes that are themselves reproducing and subject to selection. For example, in an estuary the microbial pool is likely to be comprised of a fairly constant input of upstream microbes (i.e., a fixed environmental pool, following Bruijning et al., (2022; Zeng et al., 2015, 2017)), the ‘shedding’ of microbes from hosts into the environment (Bruijning et al., 2022; Daybog & Kolodny, 2023; Zeng et al., 2015), and reproduction of microbes that are already within the environment.

Here, we develop a model framework to explore how host and microbe fitness interact with each other under selection, multiple microbial generations and mixed environmental acquisition. We model the environmental pool as comprising of a fixed environmental pool, ‘shedding’ of microbes from hosts in the environment, and – the new component we incorporate into our model - an additional environmental microbial contribution from an autochthonous source, i.e., a pool of microbes existing in the environment. We then apply this framework to understand how holobiont fitness is affected by different types of changing environments, as an oft-cited justification for holobiont research relates to microbially mediated adaptation (Baldassarre et al., 2022; Shah et al., 2024). Our model also allows for environmental variation to occur at the microbial timescale. Our work overlaps, in part, with that of Bruijning et al. (2022) (whose focus was on optimization of vertical transmission under variable environment conditions), but more realistically models the microbial environmental community, and further explores how microbial generations can offset some of the fitness costs of these challenging environments.

## Methods

### Modelling approach

Our model establishes a constant sized population (*n_h_*) of haploid hosts. A number (*N_m_*) of microbial taxa make up large microbial communities present within each host (of population size *n_mh_*) and in an external environment (of population size *n_mE_*). Both hosts and microbial taxa are assigned trait values that determine their fitness. Host trait values φ*h* are determined by both their own initial genetic trait value (*φhg)* with weighting *G*, and by the mean trait value of their microbiome *φM*, with weighting (1-*G*). The host genetic trait values are heritable and drawn from a normal distribution (V(n_H_, G._H_)). Trait values for each microbial taxon are drawn from a uniform distribution between −1 and 1. For each host generation, there are a fixed number of microbial generations both within the host and in the external environment (*T_M_* >= 1). At each generation, whether host or microbe, the communities are entirely replaced, as we do not model overlapping generations, and both hosts and microbes reproduce clonally.

The simulation begins with an environmental pool initially comprised of all microbial taxa at approximately equal abundances, and the first generation of hosts acquire their microbiome at random from this pool. The environmental pool is then regenerated each microbial generation and is made up of some proportion, *Y*, being shed from the collective host microbial pool, some proportion *Z* being made up from the previous environmental pool, and a final proportion (1-*Z*-*Y*) being made up from the fixed environment with all microbial taxa at equal abundances (Fig. 1a). Microbial taxa are sampled multinomially from each source until the total number of microbes is reached, where the probability of being sampled is the product of the fitness and the relative abundance of the microbial taxa within the relevant source. The fitness of microbial taxon *m* in the environment is calculated as:

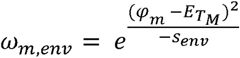

where the environmental optimum *E_T_M__* is the trait value that maximises microbe fitness in the environment at each microbial timepoint (*T_M_*) and *s_env_* > 0 is the selection parameter in the environment, with small values corresponding to strong selection. The environmental optimum is allowed to vary over time.

**Fig. 1:**
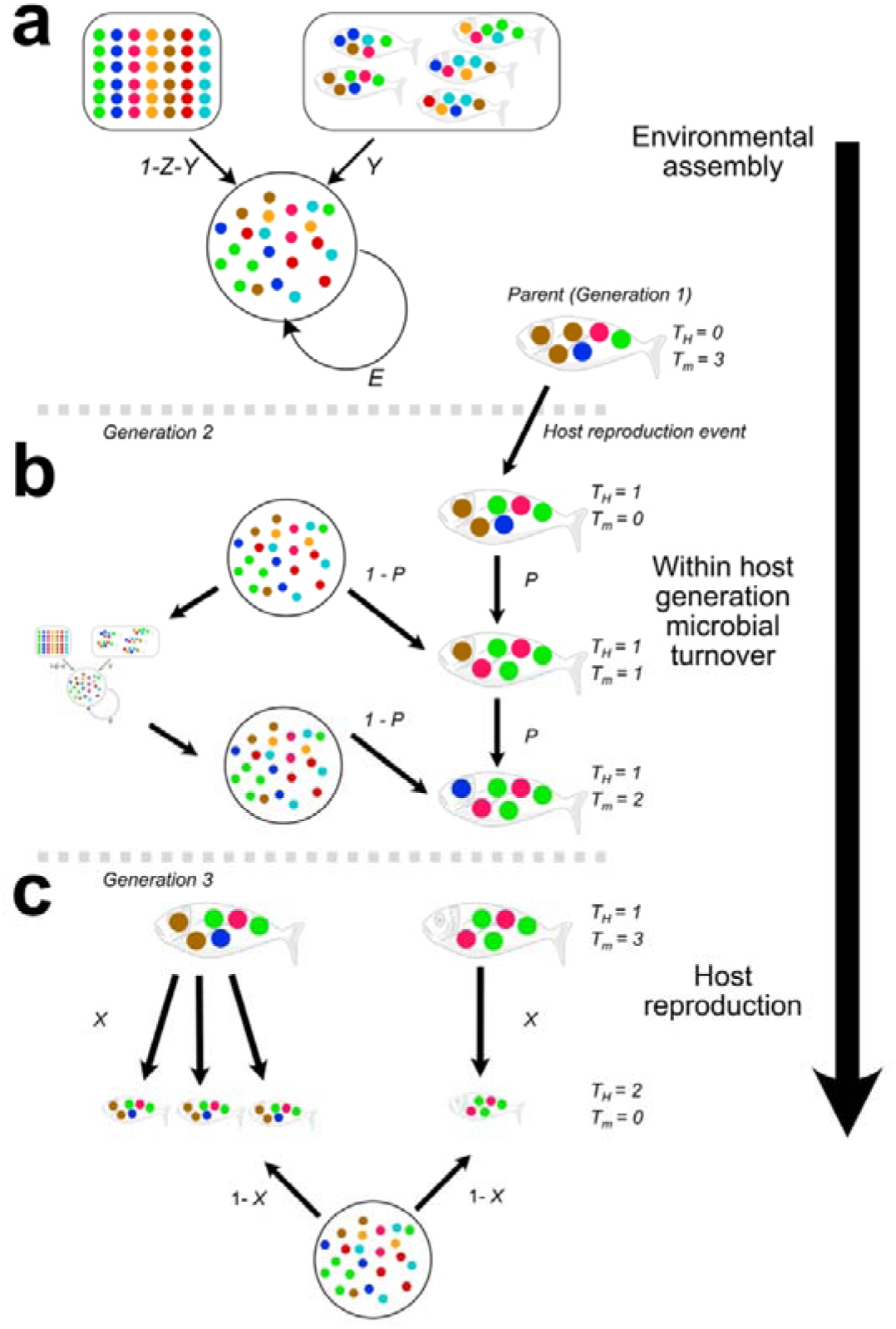
Diagram of simulation process. Arrows within each panel represent multinomial sampling events based on the product of fitness and relative abundance. In a) the environmental microbial pool is assembled based on some contribution of its previous state (*Z*), some contribution from the hosts (*Y*) and the remainder from a fixed environmental pool (1–*Z*–*Y*). b) Host microbes undergo change for a specific number of microbial generations, where the environment also undergoes change during this time. Host microbes are sampled at each microbial generation from the host with proportion *P* and from the external environment with proportion 1-*P*. c) Hosts undergo reproduction where their fitness is determined based on their microbiome, their offspring inherit proportion *X* of their microbiome from their parent, with the remainder being made up of colonizers from the environment.

Because there may be many generations of microbes within a host generation, the microbiome can change over the lifetime of the host (Fig. 1b). With each microbial generation within the host, the microbiome is sampled from its previous generation of that host at proportion *P*, and from the environment at proportion 1-*P*. As per the strategy within the environment, microbial taxa within the host are sampled multinomially from each of the now two sources, where the probability of being sampled is the product of the fitness and the relative abundance of the microbial taxa in that source. The fitness of microbial taxon *m* within a host *h* is calculated as:

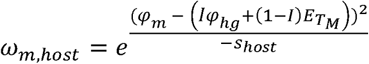

where the within-host optimum is determined by a weighting *I* of the host trait and (1-*I*) of the environmental optimum, and *s_host_* > 0 is the selection parameter in the host, with values close to 0 corresponding to strong selection, and very large values (*Ns* > 1) approaching neutrality. Note that the weightings *I* and (1-*I*) are chosen to reflect the influence of the internal host environment and external environment on a host’s microbiome.

After the specified number of microbial generations, *T_M_*, has passed, the host undergoes fitness-based reproduction (Fig. 1c), where hosts are sampled with replacement with a probability equal to their fitness. Host fitness is calculated as:

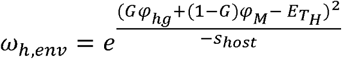

Where *E_T_H__* is the trait value that maximises host fitness in the environment at each host timepoint (*T_H_*). Following host reproduction, offspring acquire a new microbiome with some proportion, *X*, being sampled from their parent and the remainder (1-*X*) from the environment. Microbial taxa are sampled multinomially from each source until the total number of microbes is reached, where the probability of being sampled is again the product of the fitness and the relative abundance of the microbial taxa within the relevant source. A description of all parameters available in our framework, and the values used for the presented work, are available in Table 1. We also provide fully annotated code for reproducing our agent based model, along with the analyses and simulations presented herein, at https://wpearman1996.github.io/Holobiont_Bookdown/.

**Table 1:**
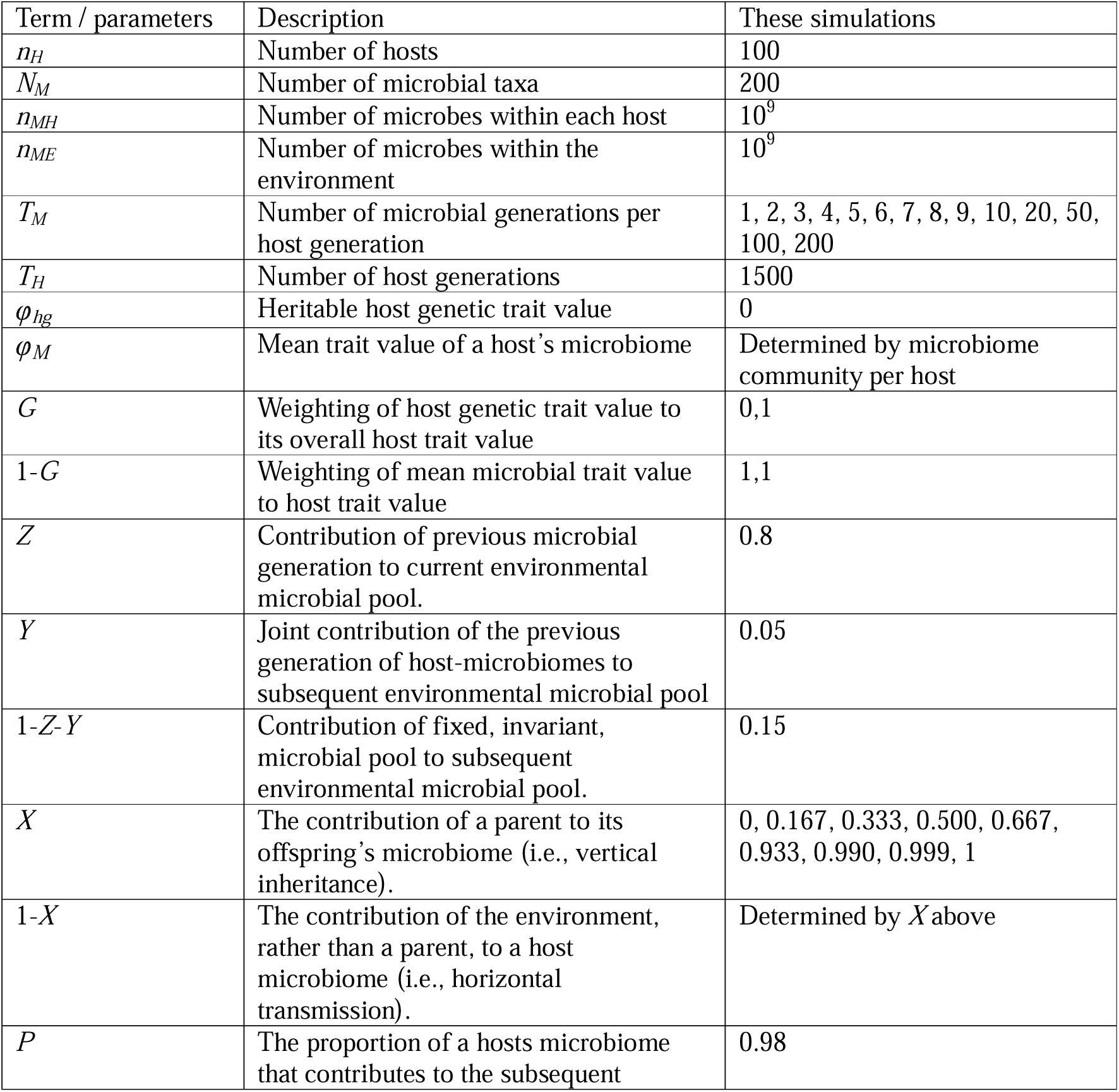
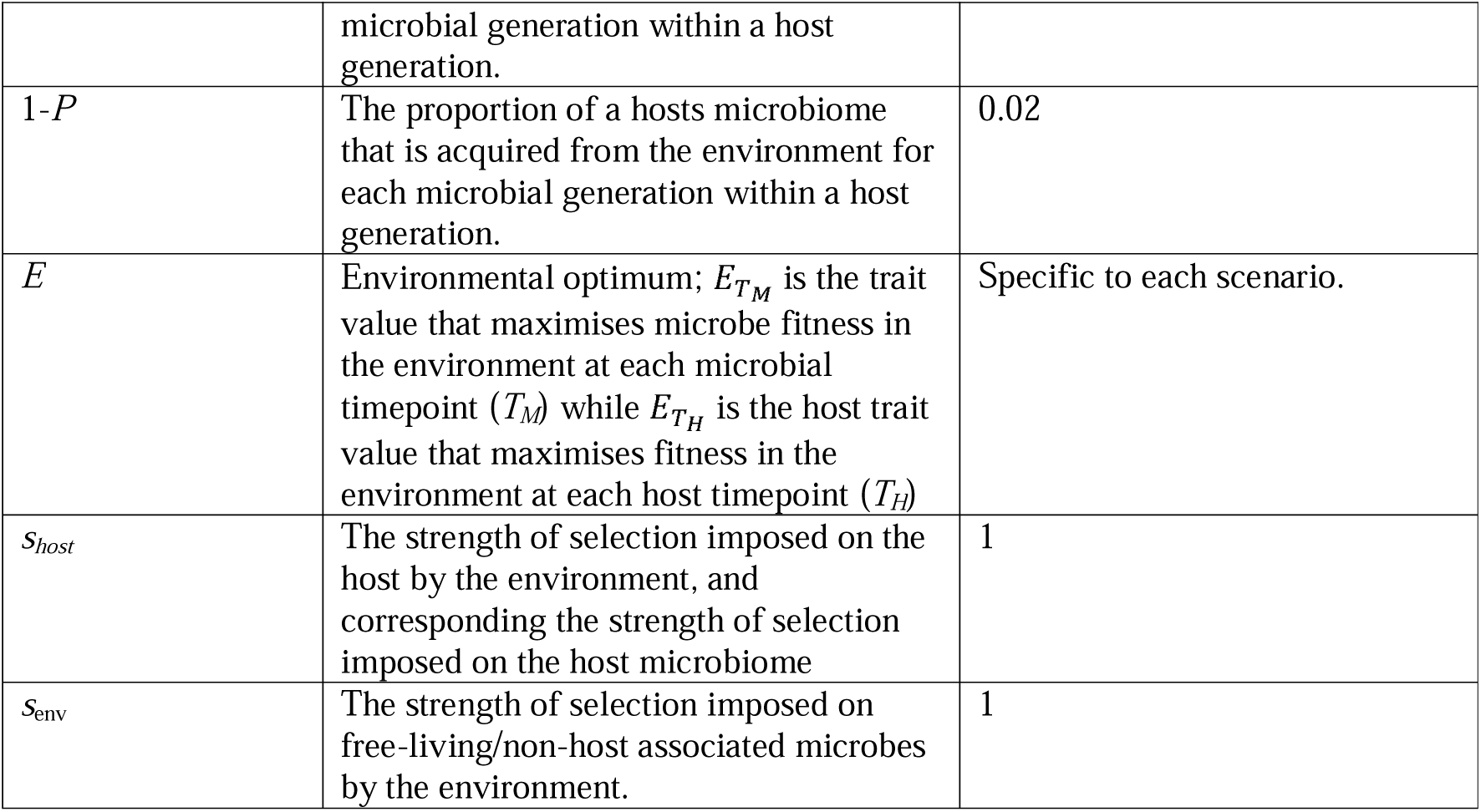
Description of parameters available in our host-microbiome modelling framework, and the choice of values specifically used in the presented simulations.

### Influence of changes in environmental conditions

We explore the influence of environmental conditions on selection in both the host and microbe, while varying the number of microbial generations per host generation (*T_M_*) and the proportion of vertical (*X*) versus horizontal inheritance (1-*X*) of microbial taxa at the point hosts reproduce. Our approach differs significantly to that of (Bruijning et al., 2022), as we simulate microbe fitness alongside host fitness rather than relying on neutral processes. Therefore, to understand how host and microbe fitness interact, we produced six scenarios of different types of changing environments exemplified in Fig. 2. In short, we produce ‘unchanging’ environments, where the environment fluctuates around a mean, an increasing mean environment where the mean environmental condition increases over time – with variation around this changing mean, and finally an environment where the variance around the mean changes over time but the mean itself does not change. Prior to implementing environmental changes for the increasing mean environment, and increasing variance environments, communities had an initial unchanging burn-in period of 200 host generations to allow for microbial communities to stabilize following initialization. For each of these scenarios we have two versions, a ‘random’ variation version where the environmental condition is drawn independently of the previous time point, or a high autocorrelation version where the environmental conditions from each point to the subsequent are highly autocorrelated.

**Fig. 2:**
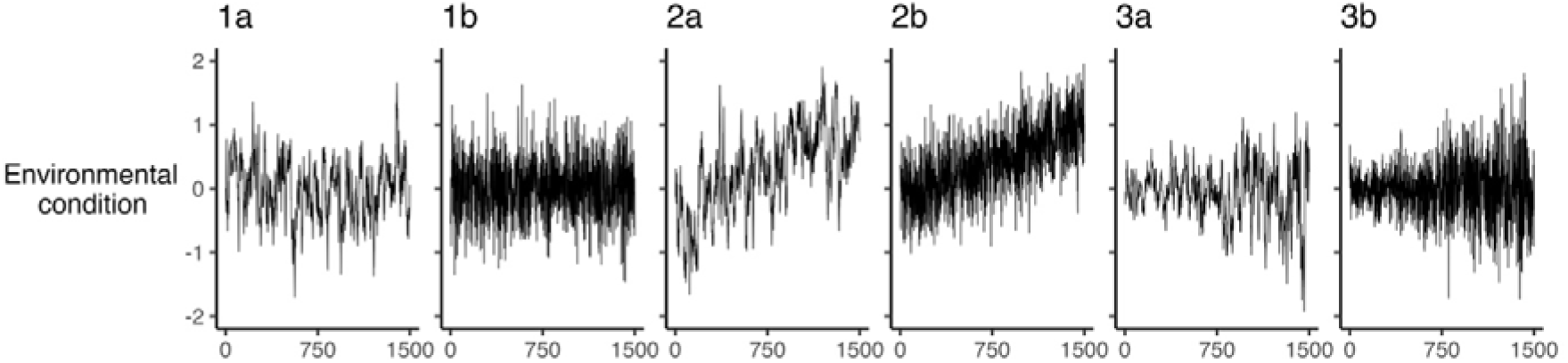
Example environmental conditions used in our simulations. Group 1 represents environmental conditions with no net change in the either mean or variance over the course of the simulation. Group 2 are environmental conditions with a net change in the mean environmental condition over the course of the simulation, but with no change in variance. Group 3 are environmental conditions where there is increasing variance, but no change in mean, over the course of the simulations. Panels “a” are those where the environmental conditions from each point to the subsequent are highly autocorrelated, while “b” are those where the environmental condition is drawn randomly and independently of the previous time point.

In our simulations, we conducted 20 replicates of each environmental condition. For each replicate the environments were generated independently – however within each replicate, we reused the same environmental conditions to produced replicate simulations where we varied specific parameters such as the number of host generations. For example, in replicate 1 of an unchanging, highly autocorrelated environment, a singular set of environmental values *E_T_H__* was generated 1,500 elements long (*T_H_*). These environmental conditions were then reused for each combination of the number of microbial generations per host generation (*T_M_*) and the proportion of vertical (*X*) versus horizontal inheritance (1-*X*). For each value of *T_M_*, the environmental conditions at *T_H_ = N* and *T_H_ = N+1* were smoothly interpolated *T_M_* times. For example, if *E_T_H__* at *T_H_ = N* is 0.2 and *T_H_ = N+1* is 0.24, and *T_M_* = 4, then the environmental conditions would be {0.2,0.21,0.22,0.23}.

Random environmental conditions were generated by drawing from a normal distribution with a mean of 0, and a standard deviation of either 0.5, or a standard deviation increasing from 0.3 to 0.85 over the course of the simulation. In the case of the high autocorrelation environment, values were drawn using a bespoke autoregressive integrated moving average (ARIMA) function to allow for varying standard deviations with an AR component of 0.9, MA component of 0, and with standard deviations varying from 0.1 to 0.3 – these values were chosen to match to match the overall standard deviations across random and high autocorrelation environmental conditions.

Our simulations covered a range of microbial generations per host generation (*T_M_* = 1, 2, 3, 4, 5, 6, 7, 8, 9, 10, 20, 50, 100, 200) and a range of vertical inheritance at host reproduction (*X* = 0, 0.167, 0.333, 0.500, 0.667, 0.933, 0.990, 0.999, 1). In all simulations, a total of *N_h_* = 100 hosts each have a microbial population of *n_M_* = 10^9^ drawn from *N_M_* = 200 microbial taxa. Host genetic trait values are set to 0 and are assigned to all members of the host population such that they are all haploid clones. The overall host trait value is determined solely by its microbial population (*G* = 0). All selection parameters were set to 1. We set *I* = 0.5 such that both the internal host genetic environment and the external environment have equal weighting on the fitness of microbes within the host. We set *P* = 0.98 such that 2% of microbes within the host are acquired externally from the environment compared to the previous microbial pool in the host. Finally, *Z* = 0.8 and *Y* = 0.05 were set to establish the contributions of the current environmental microbial community, host shedding of microbes, and a fixed microbial input to the environmental microbial community (also consisting of 10^9^ microbes).

When a host reproduces, microbes are inherited at proportion *X* from parent to offspring, but subsequently at proportion *P* from one microbial generation within the host to the next. We generally expect that *X* and *P* will differ, because colonization of a new host is likely to differ from the microbial turnover in an existing host microbial community. Further, the impact of the initial colonization of a new host will become diluted as the number of microbial generations within the host increases. We therefore tracked the effective vertical inheritance from one host generation to the next as the proportion of the original colonization community that was remaining at the end of the host generation. Vertically acquired microbes were classified as any microbe originating from the host’s direct parent, thus if a parent sheds a microbe into the environment – which then persists for a few microbial generations before colonizing the offspring – this would also be classified as vertical inheritance. We quantified effective vertical inheritance independently for each environmental scenario (Fig. 2) at the 500^th^ host generation, following this we ran an additional set of simulations for *T_M_* = 1, where *X* was set as the effective vertical inheritance equivalent as quantified for *T_M_* = 200.

For each simulation, at the end of every host generation (i.e., prior to reproduction) we recorded the average host fitness and average microbial fitness within the host, as defined by the equations above. Further, we recorded average microbial alpha diversity across hosts for each replicate host as the scaled Shannon-Weiner index following Zeng et al. (2015).

## Results

We first present case studies where *T_M_* = 1 and *T_M_* = 200 across different environments and levels of vertical inheritance in order to describe changes in fitness and diversity over the course of the simulation. We then present fitness and diversity results at the end of each simulation, so that the impact on these metrics across the range of values of *T_M_* can be explored. Finally, we explore the role of effective vertical inheritance in shaping fitness.

### Dynamics of microbe and host fitness where *T_M_* = 1

We first explored the changes in microbial and host fitness over time across environmental conditions when there was only one microbial generation per host generation. Beyond the initial stabilization of the communities in the first generations, microbial fitness was largely unchanging in a variable environment that does not change systematically over time, whether environmental conditions were random or had high autocorrelation (Fig. 3a-b). In contrast, host fitness declined towards the end of these simulations under random environmental conditions (Fig. 3h), but not for an autocorrelated environment (Fig. 3g). We find that under environments with increasing variance over time, microbial fitness decreases in both random and high autocorrelation environments (Fig. 3e-f). With a net environmental change over time but no change in variance, microbial fitness decreases only when there is complete vertical inheritance of the microbiome, but remains steady otherwise (Fig. 3c-d). Conversely, host fitness declines over time both with changing environmental mean and with increasing environmental variance (Fig. j-k). Under complete vertical inheritance, there was a notably larger decrease in host fitness under changing environments experiencing high autocorrelation (Fig. 3j) but not for those changing at random (Fig. 3h), this decrease in fitness was similar to when the microbiome contributed nothing to host fitness (black line, Fig. 3g-k), while in all other instances host fitness derived from the microbiome was either larger or similar than when fitness was derived from the host’s own trait value.

**Fig. 3:**
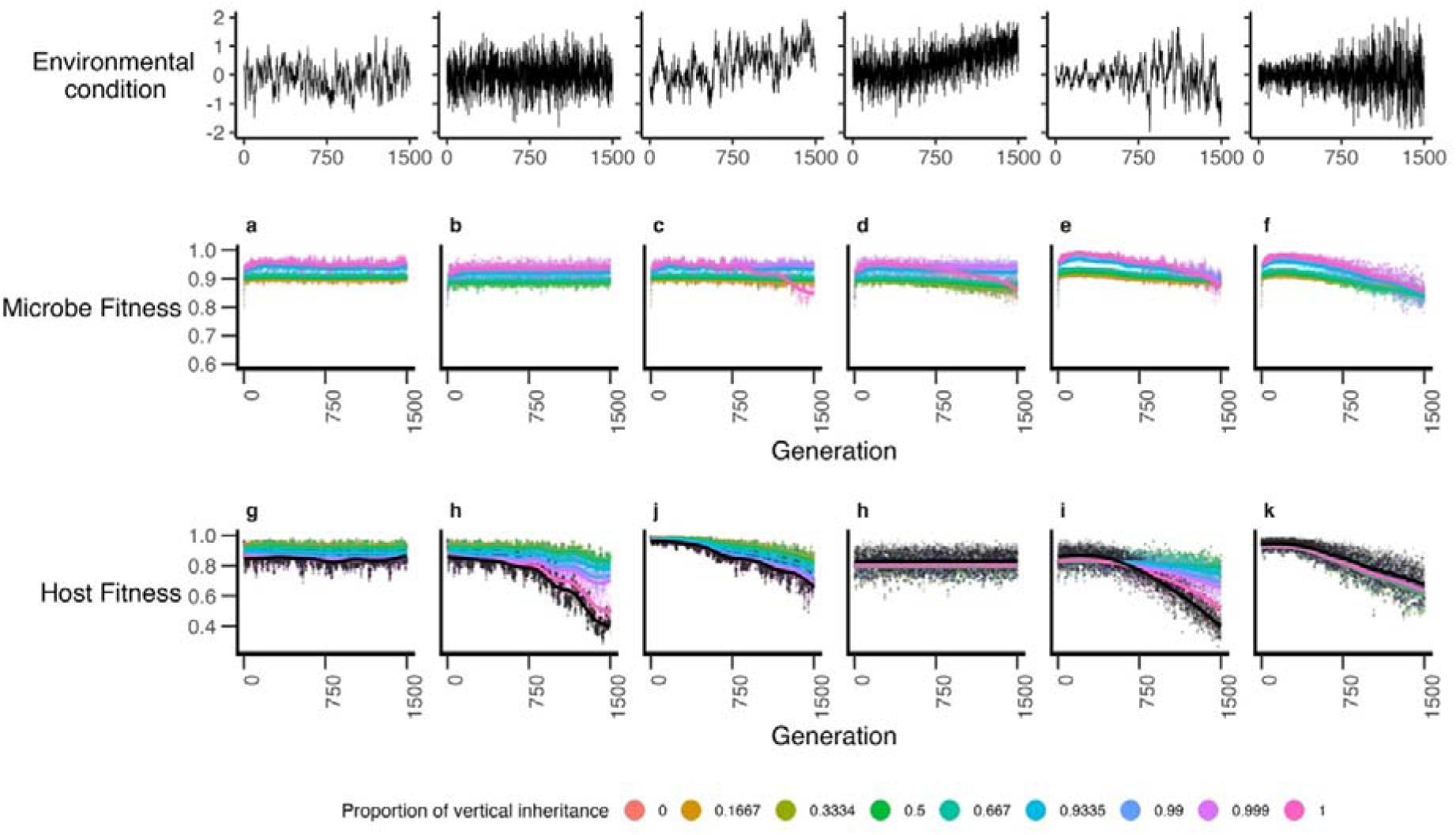
Microbial fitness (a-f) and host fitness (g-l) over the course of the simulations when the number of microbial generations per host generation is 1. Lines represent GAM smoothing; data represents mean values from 20 replicate simulations. Black points in panels g-k are the host fitness where the contribution of the microbiome to host fitness is zero, this value does not affect microbial fitness and as such is not displayed on a-f.

In addition to the relationships with environmental conditions, we find that host and microbial fitnesses were regularly at odds with each other, where the degree of vertical inheritance which maximizes their respective fitness were in conflict between host and microbe. More explicitly, high levels of vertical inheritance appear to lead to higher microbial fitness, but lower host fitness. We found that complete vertical inheritance when *T_M_* = 1, tended to result in equivalent fitness to a simulation where hosts derived their fitness entirely from their own genotype rather than entirely from the microbiome. Beyond this, we found that any level of vertical inheritance < 1 resulted in higher host fitness (Fig. 3g-k).

### Dynamics of microbe and host Fitness – *T_M_* = 200

When the number of microbial generations per host generation (*T_M_*) was increased to 200, we observed that in all instances fitness of both host and microbe increased relative to *T_M_* = 1 (Fig. 3 & 4). We found that increasing *T_M_* led to a less severe decline in both host and microbial fitness, and lead to the homogenization of the influence of vertical inheritance with regards to host fitness (Fig. 4g-k). In other words, at high values of *T_M_* there was no impact of any level of vertical inheritance on host fitness regardless of environmental conditions (Fig. 4). We found that microbially-derived host fitness was typically much higher than when host fitness was entirely derived from its own genetic trait value (Fig. 4g-k). At the same time, microbial fitness remained highest under higher levels of vertical inheritance (Fig. 4a-f). As with the *T_M_* = 1 scenario, we found that when *T_M_* = 200 host fitness was always greater when derived from the microbiome than from the host’s own trait value (Fig. 4g-k).

**Fig. 4:**
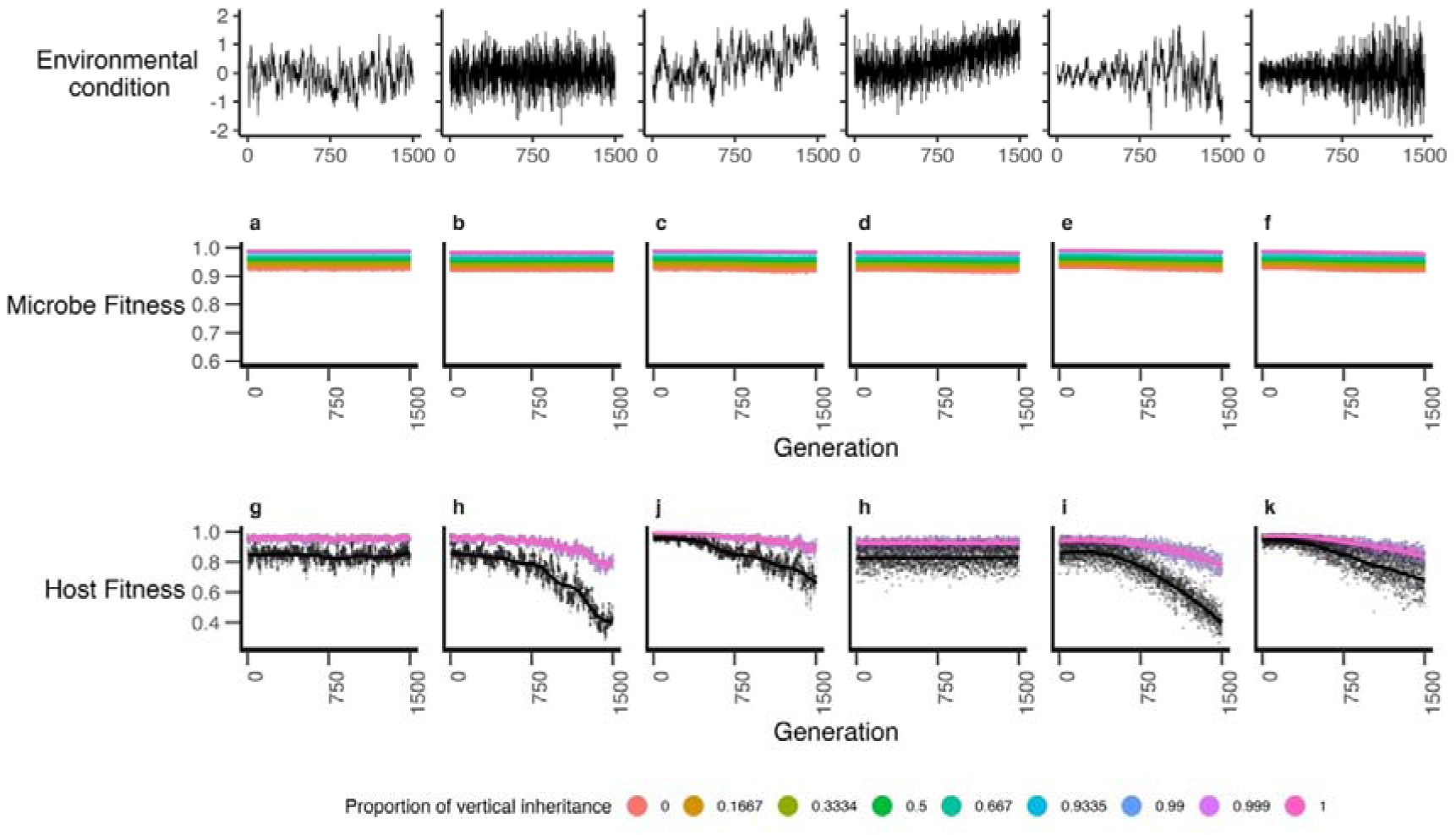
Microbial fitness (a-f) and host fitness (g-l) over the course of the simulations when the number of microbial generations per host generation is 200. Lines represent GAM smoothing; data represents mean values from 20 replicate simulations. Black points in panels g-k are the host fitness where the contribution of the microbiome to host fitness is zero, this value does not affect microbial fitness and as such is not displayed of a-f.

### Dynamics of diversity

Over time when the number of microbial generations per host generation (*T_M_*) was 1, alpha diversity initially declined at higher levels of vertical inheritance (*X*) followed by a plateau (Fig. 5-a-f). Most notably, when *X* = 1 diversity declined to practically zero. Conversely, when *T_M_* = 200 diversity largely remained static across the environmental conditions (Fig. 5 g-l). Broadly, low values of vertical inheritance resulted in higher levels of alpha diversity regardless of environmental conditions.

**Fig. 5:**
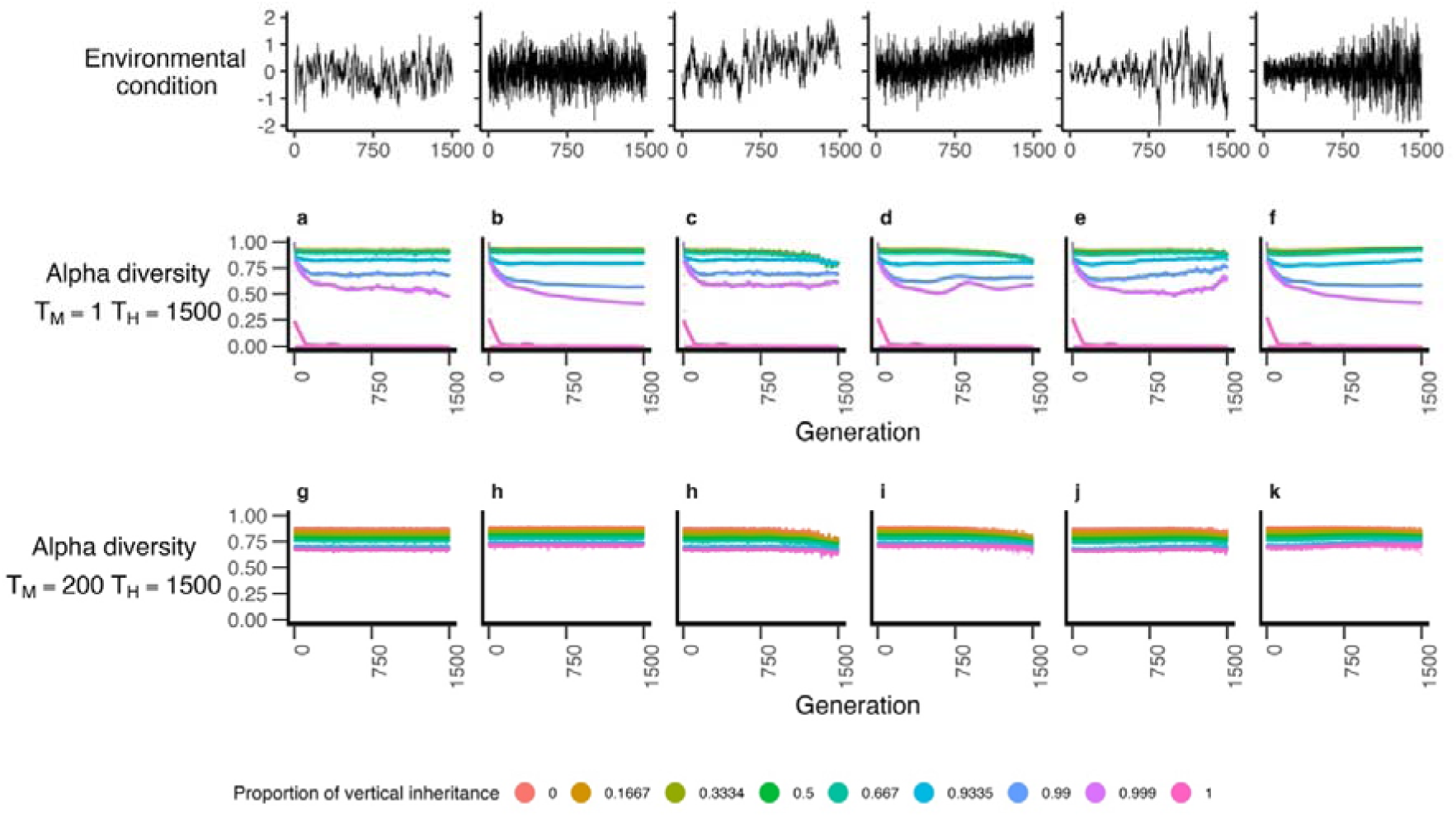
Alpha diversity of microbiomes at host generation 1,500 in response to both vertical inheritance proportion and number of microbial generations per host generation (a-f), alpha diversity over the course of the simulations when the number of microbial generations per host generation is 1 and (g-k) over the course of the simulations when the number of microbial generations per host generation is 200. Lines represent GAM smoothing; data represents mean values from 20 replicate simulations.

### Influence of *T_M_* on final fitness

Generally, we found that higher numbers of microbial generations per host generation (larger *T_M_*) lead to higher fitness of both hosts and microbes. When there was a net change in the environmental conditions over the course of the simulation, complete vertical inheritance in tandem with *T_M_* = 1 resulted in substantially lower fitness than other scenarios (Fig. 6 c-d, h-i). These net changing environmental conditions also resulted in host fitness being less influenced by higher values of *T_M_* and in fact were marginally higher at low levels of vertical inheritance and *T_M_* (Fig. 6h-i).

**Fig. 6:**
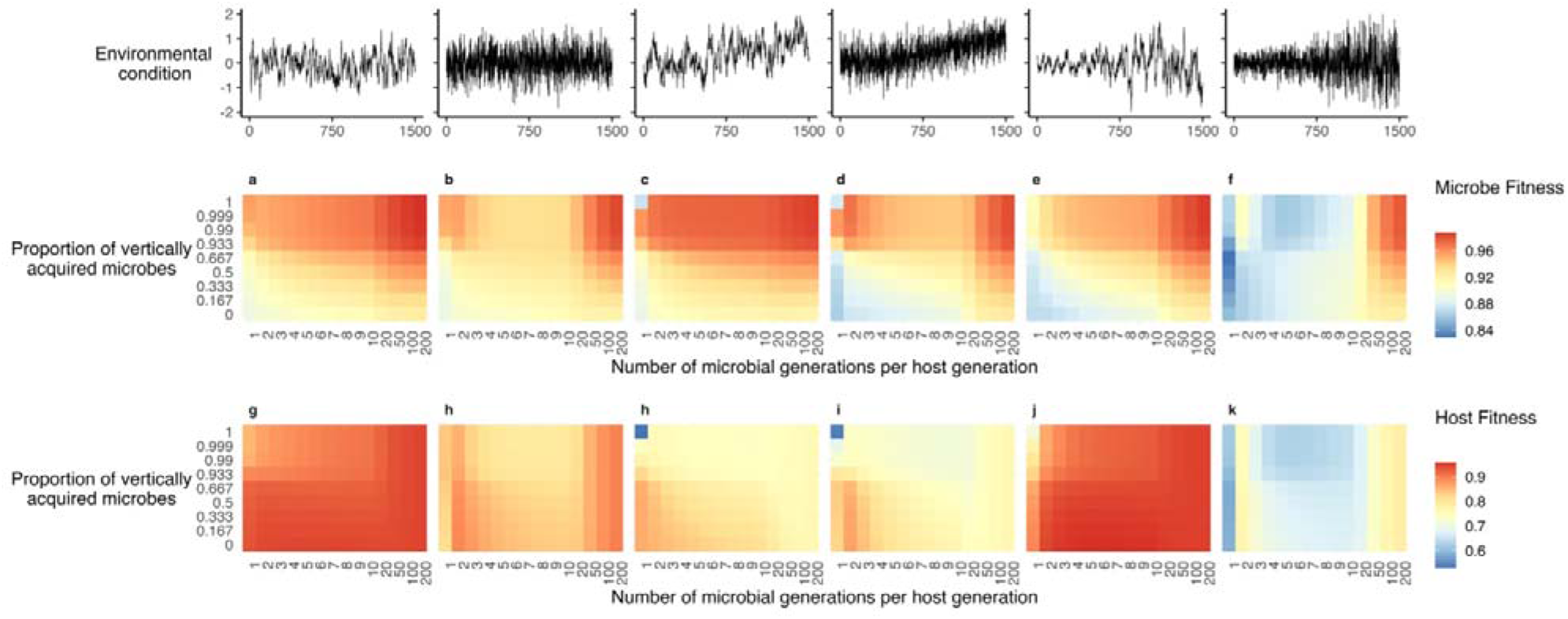
Microbe fitness at host generation 1,500 in response to both vertical inheritance proportion and number of microbial generations per host generation (a-f), host fitness at generation 1,500 (g-l) in response to both vertical inheritance proportion and number of microbial generations per host generation.

### Influence of *T_M_* on final diversity

We found that low values of either vertical inheritance or *T_M_* maximized alpha diversity (Fig. 7), while high values of *T_M_* resulted in marginally lower diversity (Fig. 7). At the same, net changing environmental conditions (Fig. 7c-d) tended to have lower diversity than other scenarios. Finally, high values of vertical inheritance where *T_M_* = 1 resulted in the lowest diversity across all scenarios (Fig. 7).

**Fig. 7:**
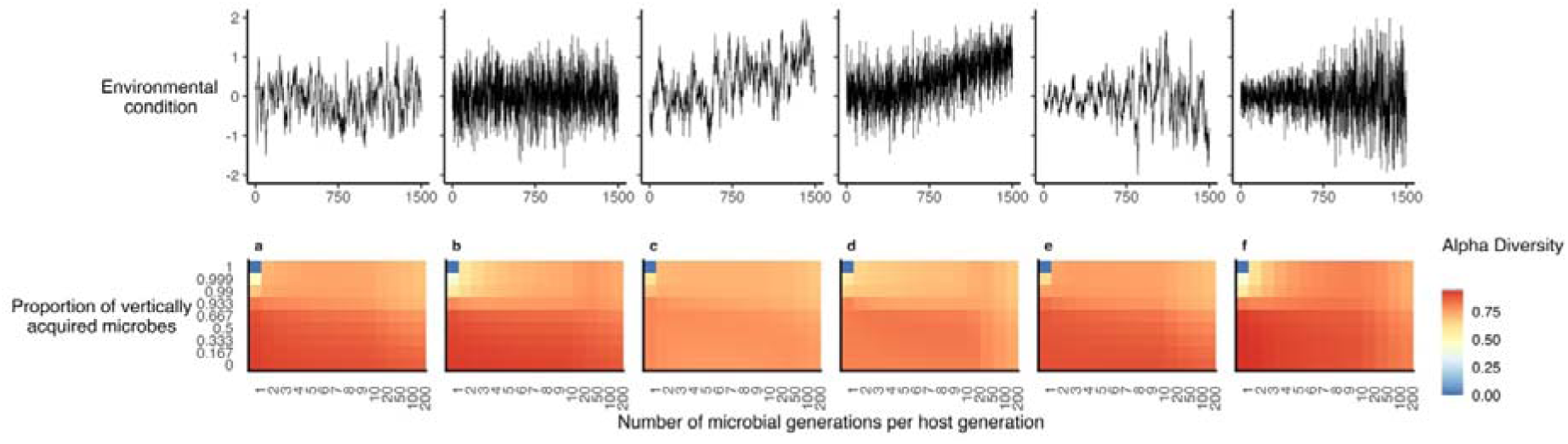
Alpha diversity of microbiomes at host generation 1,500 in response to both vertical inheritance proportion number of microbial generations per host generation (a-f), alpha diversity over the course over time(g-l) over the course of the simulations when the number of microbial generations per host generation is 200. Lines represent GAM smoothing, data represents mean values from 20 replicate simulations.

To ascertain the importance of vertical inheritance independently of microbial generation count per host generation, we quantified effective vertical inheritance for each value of *X* under each environmental scenario, across a range of values of *T_M_*. We then ran the simulation for *T_M_* = 1 at these lower effective vertical inheritance values, and compared host and microbe fitness to their fitness at *T_M_* = 200. We show that both host and microbe fitness overall increase with increasing numbers of microbial generations, in agreement with our previous conclusions. These results suggest that high values of *T_M_* result in fitness gains regardless of the level of effective vertical inheritance (Fig. 8b-m).

**Fig. 8:**
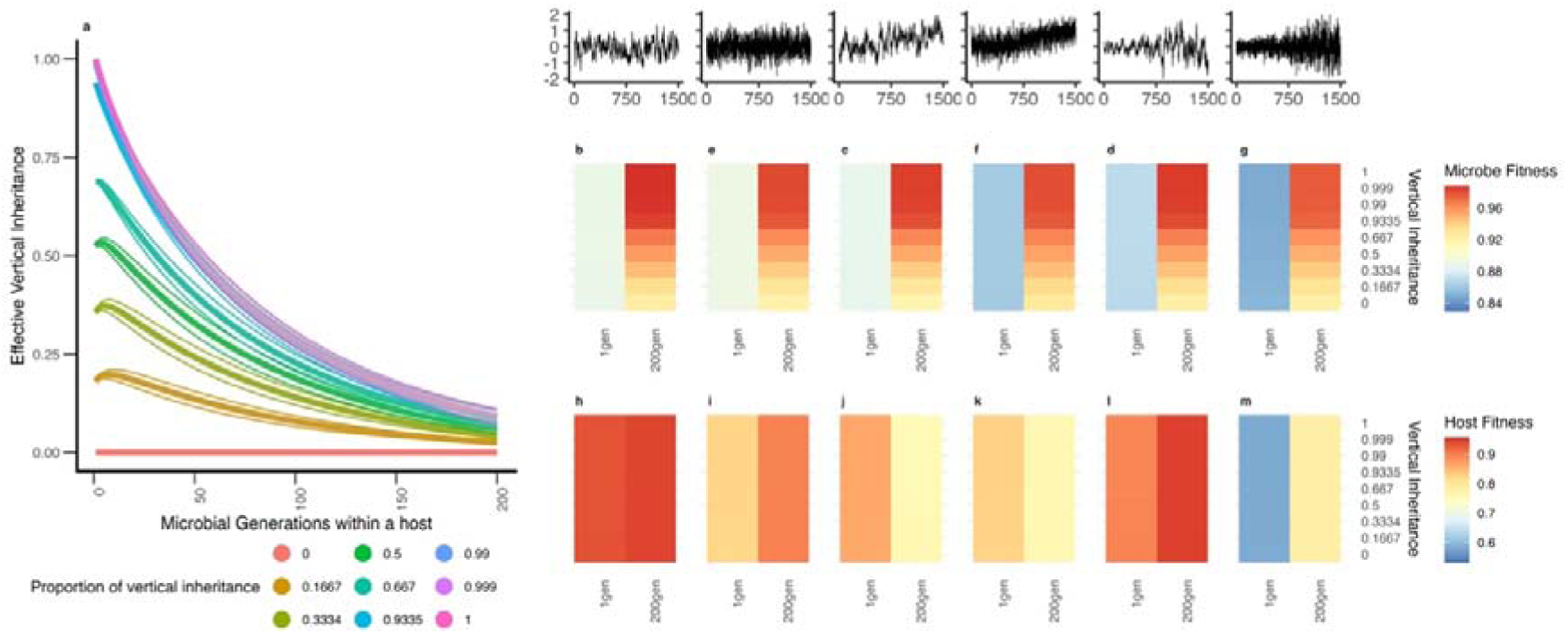
Patterns of decay in effective vertical inheritance within a host generation for varying levels of initial vertical acquisition (a). Microbe (b-g) and host (h-m) fitness at host generation 1,500 for 200 microbial generations per host generation for effective vertical inheritance at different levels. The effective vertical inheritance is standardized such that “1 gen” was run with effective vertical inheritance of 200 generations found in (a). For example, if vertical inheritance is complete but vertical inheritance at 200 generations was determined as 10%, then we initialized a simulation with a vertical inheritance of 10% for 1 generation – and compared this to a simulation where we allowed the simulation to progress for 200 microbial generations where we initialise the vertical inheritance as 100%.

## Discussion

Understanding the formation and eco-evolutionary processes of holobionts is a key objective in the study of host-associated microbiomes (Leonard et al., 2024). Using simulations, we find that multiple microbial generations per host generation is a key factor that shapes the fitness of simulated holobionts. Most studies to date have made key simplifications when simulating host microbiome evolution – assuming either neutrally assembled microbial communities while having non-neutrally assembled host communities, or restricting simulations to one microbial generation per host generation. We show here that consideration of both factors is important to understanding how host-microbe systems can evolve under varying environmental conditions.

### Diversity

Our results are somewhat intuitive with regards to alpha diversity and support those of Zeng et al., (2015 & 2017). First, under complete vertical inheritance (*X* = 1) and when the number of microbial generations per host generation (*T_M_*) is extremely low, alpha diversity declines rapidly to virtually zero (Fig 5a-f). This occurs because there are very few opportunities for a host to gain new microbes – thus when a microbe is lost, it is effectively lost permanently and so diversity can only decline. More generally, high levels of vertical inheritance limit the number of microbial migrants from the environments at birth and result in low levels of alpha diversity. We find that the highest alpha diversity is therefore observed at both low levels of vertical inheritance and high numbers of microbial generations per host generation, as environmental colonisation results in greater turnover and replacement of the microbiome both within and across each host generation – something also observed by Zeng et al. (2015).

In our simulations, the environment experienced by the microbiome is a smooth interpolation of the environment conditions at each host generation. For microbial communities with multiple generations per host generation, this approach creates a continuous transitioning environment, and thus generally promotes higher microbial diversity relative to fewer microbial generations per host generation. We note that there is typically higher diversity in the autocorrelated environmental conditions relative to the randomly generated counterpart. This follows from the Intermediate Stochasticity Hypothesis, which posits that intermediate levels of environmental variability can maximize diversity through temporal niche differentiation – where under static environments one or few specialists can dominate, and under highly changeable environments nothing or few things can survive (Santillan & Wuertz, 2022). Similar results have also been observed in macroalgal microbiomes, where environmental predictability (as measured through the degree of temporal autocorrelation in temperature) is a driver of holobiont diversity (Pearman et al., 2024).

### Fitness

First, we demonstrate that host fitness derived from the microbiome was almost always greater than fitness derived from a host’s own trait value. Although expected, this result demonstrates that the microbiome can be important in buffering hosts to environmental change. We found both host and microbe fitness increased with higher numbers of microbial generations per host generation (i.e. *T_M_*), such a result makes sense because high values of *T_M_* ‘spread’ the environmental change over more microbial generations, as described above. Such ‘spread’ allows for gradual adjustment and changes in the microbiome, whereas when environments undergo sudden drastic changes, this leads to a stronger disturbance effect and thus greater initial changes in the microbiome. In the host, microbe fitness is acted upon both by the host and the environment and thus higher numbers of microbial generations enable the microbial community time to change in response to this joint selection, to the benefit of both host and microbiome fitness.

Interestingly, broadly across our simulations we found that low values of *T_M_* resulted in a conflict between the influence of vertical inheritance on host and microbe fitness, where high vertical inheritance maximized microbial fitness within the host but tended to have the opposite effect on host fitness. This result highlights the contrast, and potential conflicts, that can arise between hosts and their microbes. Importantly, because we model host fitness as derived from the relationship between the average trait value of a host’s microbial community and its distance to the environmental condition, it is best for a host to completely replace its microbiome when *T_M_* is low so that it can acquire a microbiome that is more environmentally fit. Notably, this does not provide benefits for the microbiome, which is suited to the environment more than the host, but allows the host to maximize the gains from the microbiome. In contrast, under low *T_M_* microbial fitness is highest under high vertical transmission because there is continuity in the microbial communities across host generations, allowing for microbial selection to operate across multiple host generations and therefore allowing for host microbiomes to adapt to the host.

A higher number of microbial generations per host generation increases both host and microbe fitness, and we hypothesise above that this is because the rate of environmental change that microbes are subject to is slowed, such that a smooth community transition can occur within a host generation, rather than undergoing major disturbances which may lead to the loss of retained microbes. The retention of these microbes leads to cross-generational priority effects which, in our simulations, may provide a benefit to hosts if those microbes subsequently become beneficial when environmental conditions change. For example, take a scenario where the environmental conditions transition from 0.2 to 0 and back to 0.2 over two host generations – if vertical inheritance is sufficiently high, some of those microbes suited to an environment of 0.2 may persist throughout the environmental change, resulting in the hosts deriving a benefit from the priority effects when the environment returns to 0.2. This possibility is further amplified by having very large microbial communities, resulting in a greater number of ‘persister’ microbes which have the advantage of not needing to recolonize the host. These cross generational priority effects appear to be stronger in our autocorrelated environments, which tend to have both higher host and microbe fitness (Fig. 5) – this is likely because autocorrelation increases the ‘continuity’ of environmental conditions, and leads to weaker disturbance effect as a result of less sharp changes in selective pressures imposed by the environment. This is perhaps also why we typically observe higher phenotypic variance (phenotype is measured as the average microbial trait value of a host microbiome, upon which selection on hosts acts) under autocorrelated environmental conditions when the number of microbial generations is low, as these cross generational priority effects enable maintenance of higher levels of diversity upon which selection can act (Supp Figs. 1-2). Such cross-generational priority effects may also be why host fitness tended to decline in an auto-correlated environment, while remaining steady in a randomly generated one. Our results are also in line with those observed by Bruijning et al (2022; Fig. 6), who found highest phenotypic variance in lower vertical transmission/high microbial generation count scenarios – although notably our results are dependent on the specific environmental conditions being studied (Supp. Figs. 1-2).

However, it is possible that the host and microbe fitness gains we see from increasing *T_M_* are primarily driven by changes in effective vertical inheritance. Effective vertical inheritance is the term we use to describe the proportion of parentally acquired microbes remaining in a host microbiome at the point at which that host reproduces. Such a value is expected to decline over time due to the loss of incumbent microbes and replacement with microbes from the environment (see microbiome heritability; Bruijning et al. 2020). Thus, even when a host acquires their initial microbiome entirely from their parents, after 100 generations only ∼10% of the parentally acquired microbes may have any direct descendants within the final host microbiome. To test the argument that effective vertical inheritance drives holobiont fitness rather than microbial generation count, we simulated equivalent effective vertical inheritance scenarios, but one where *T_M_* = 1 and the other where *T_M_* = 200. Our results demonstrate that there is usually higher fitness in both host and microbe in the latter, supporting our hypothesis that the number of microbial generations per host generation acts as a buffering effect to environmental change.

Given that optimisation of the degree of vertical transmission is important to maximizing host fitness under variable environments (Bruijning et al., 2022), it is possible that effective vertical inheritance may offset some of the fitness benefits of being long-lived. For example, a host which has 200 generations of microbes per generation may have an effective vertical inheritance value of 11%, despite an initial vertical inheritance value of 100% - thus a shorter-lived organism with the same effective vertical inheritance may be equally fit. This suggestion is further reinforced by work by van Vliet & Doebeli (2019), who suggest that long-lived organisms, or organisms with which there is a high decay in parentally acquired microbes over time, are less likely to evolve as a ‘holobiont’. The reason for this suggestion is based on the argument that enough parental microbes must be present at the point of reproduction to enable holobionts to evolve in a neo-Darwinian fashion – although this approach differs from our model, in that it assumes that it is costly for a ‘holobiotic’ microbe to be associated with a host. In contrast to this expectation, we observe that both host and microbe fitness are typically maximized at higher values of T_M_ and vertical inheritance – such a scenario leads to a maximization of effective vertical inheritance while preserving the benefits gained by the responsiveness of the microbiome to environmental pressures through environmental acquisition of microbes.

One reason that we suggest higher generational counts is favourable is due to the impact of sudden disturbances on microbial communities. For example, when a community undergoes an instantaneous major environmental change (such as at *T_M_* = 1 with a randomly generated environment) – this could lead to a loss of much of the diversity in the microbiome and aligns with research into mammalian microbiomes, where post-disturbance alpha diversity is 43% of the pre-disturbance community (Jurburg et al., 2024). However, because we smoothly interpolate between subsequent environmental values for microbial generations there will be a smooth transition between host generations when *T_M_* = 200 – reducing the impact of the disturbance. These gradual changes in diversity can minimize the invasion of microbes into the community through density blocking and priority effects. Furthermore, even when disturbances are strong enough to have an impact in *T_M_* = 200 scenarios, the high generation count provides a recovery phase to communities. Once again, these results align with those of mammalian microbiomes which exhibit recovery of almost all pre-disturbance diversity within ∼50 days post-disturbance (Jurburg et al., 2024).

### Outlook and next steps

By extending existing models of microbiome assemblages, we have made significant contributions to our understanding of how microbial generation count and changing environments impact both host and microbe fitness in simulated holobionts. However, there is a clear need to further extend this and other existing models in order to understand how microbiome assemblages can evolve, and may buffer hosts from the impacts of a changing environment. Perhaps one of the most pressing elements to focus on is the genetic elements of the host. For example, many studies to date, including this one, have made the key assumption that a host’s fitness is entirely derived from its microbiome. This is a necessary simplification, because it sidesteps the need to develop a quantitative genetic model (inclusive of mutation) to describe host evolution within a holobiont system. Once developed, and as previously suggested by Daybog & Kolodny (2023), an exciting further extension of such a host genetic model would be to allow the contribution of the microbiome to vary across hosts and evolve over time, to test the conditions under which host-microbiome partnerships, i.e., holobionts, are most likely to evolve.

## Code Availability

Code for reproducing our analyses and simulations, and example code to demonstrate the model is available at: https://wpearman1996.github.io/Holobiont_Bookdown/. *An archived version of the code will be deposited into zenodo with a DOI upon acceptance of the manuscript*.

## Acknowledgements

We would like to thank members of the Santure lab for helpful discussions around both designing and troubleshooting the simulations. We thank Marti Anderson for helpful discussions that guided the manuscript. We thank Qinglong Zeng and Yao Xiao for their helpful guidance on the implementation and design of other modelling approaches. We also wish to acknowledge the use of New Zealand eScience Infrastructure (NeSI) high performance computing facilities, consulting support and/or training services as part of this research. New Zealand’s national facilities are provided by NeSI and funded jointly by NeSI’s collaborator institutions and through the Ministry of Business, Innovation & Employment’s Research Infrastructure programme. URL https://www.nesi.org.nz.

## Supplementary Material

**Supp. Fig. 1:**
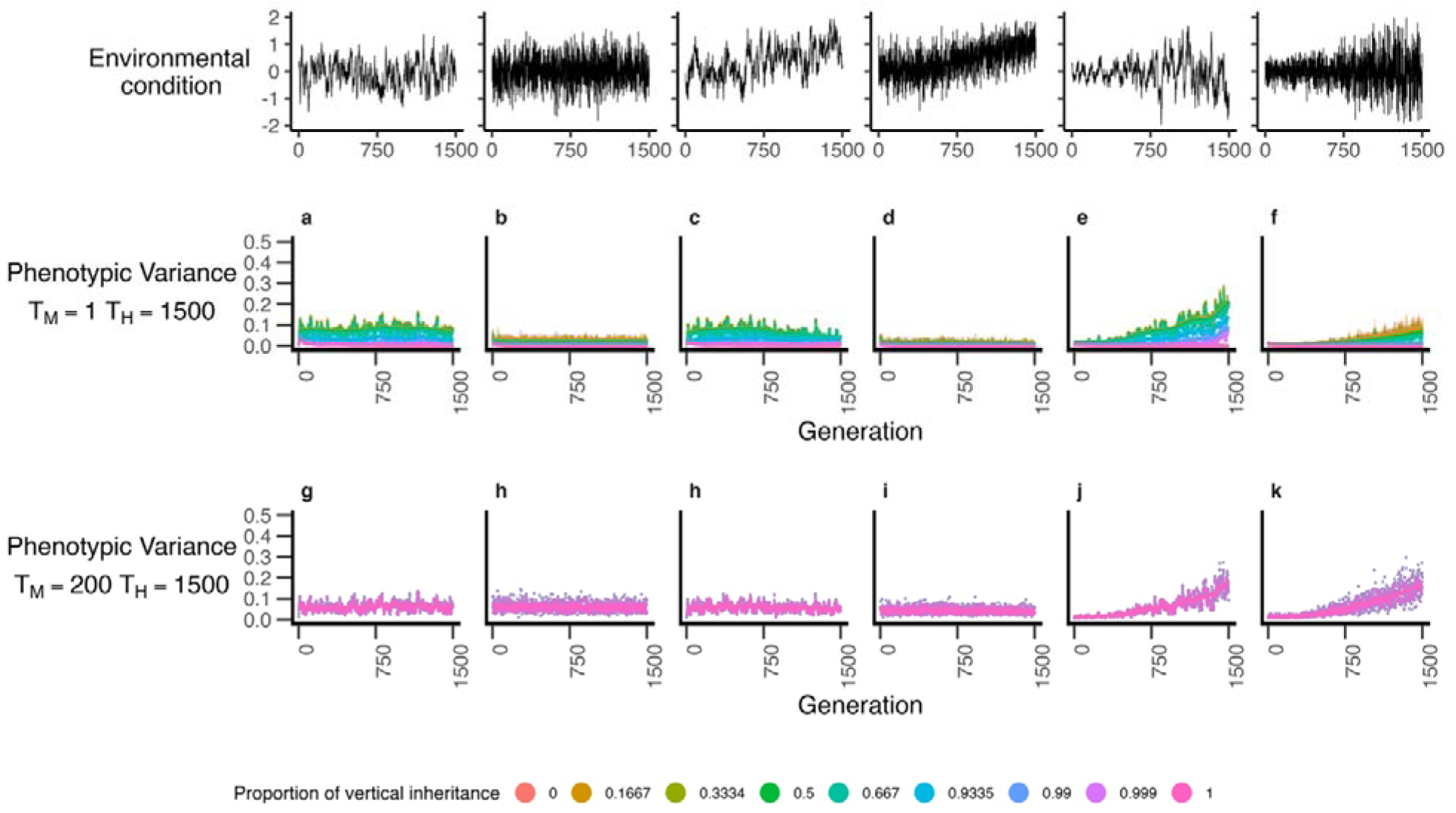
Phenotypic variance of hosts over the course of our simulations where the number of host generations is 1(a-f) or is 200 (g-k). Lines represent GAM smoothing; data represents mean values from 20 replicate simulations.

**Supp. Fig. 2:**
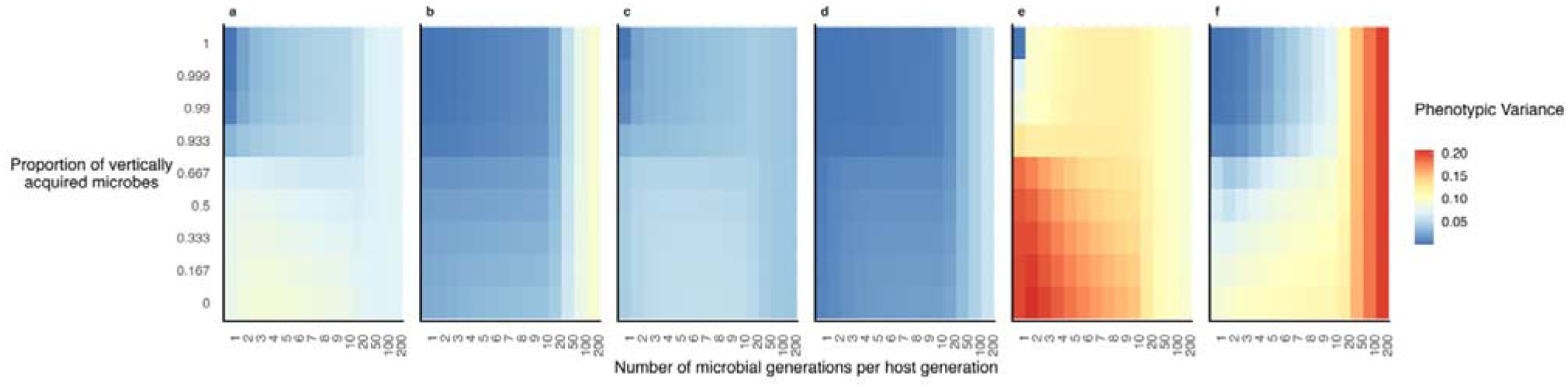
Phenotypic variance of hosts at generation 1,500 in response to both vertical inheritance proportion and number of microbial generations per host generation (a-f).

