## Supplementary material for "Joint consideration of selection and microbial generation count provides unique insights into evolutionary and ecological dynamics of holobionts": Supp.

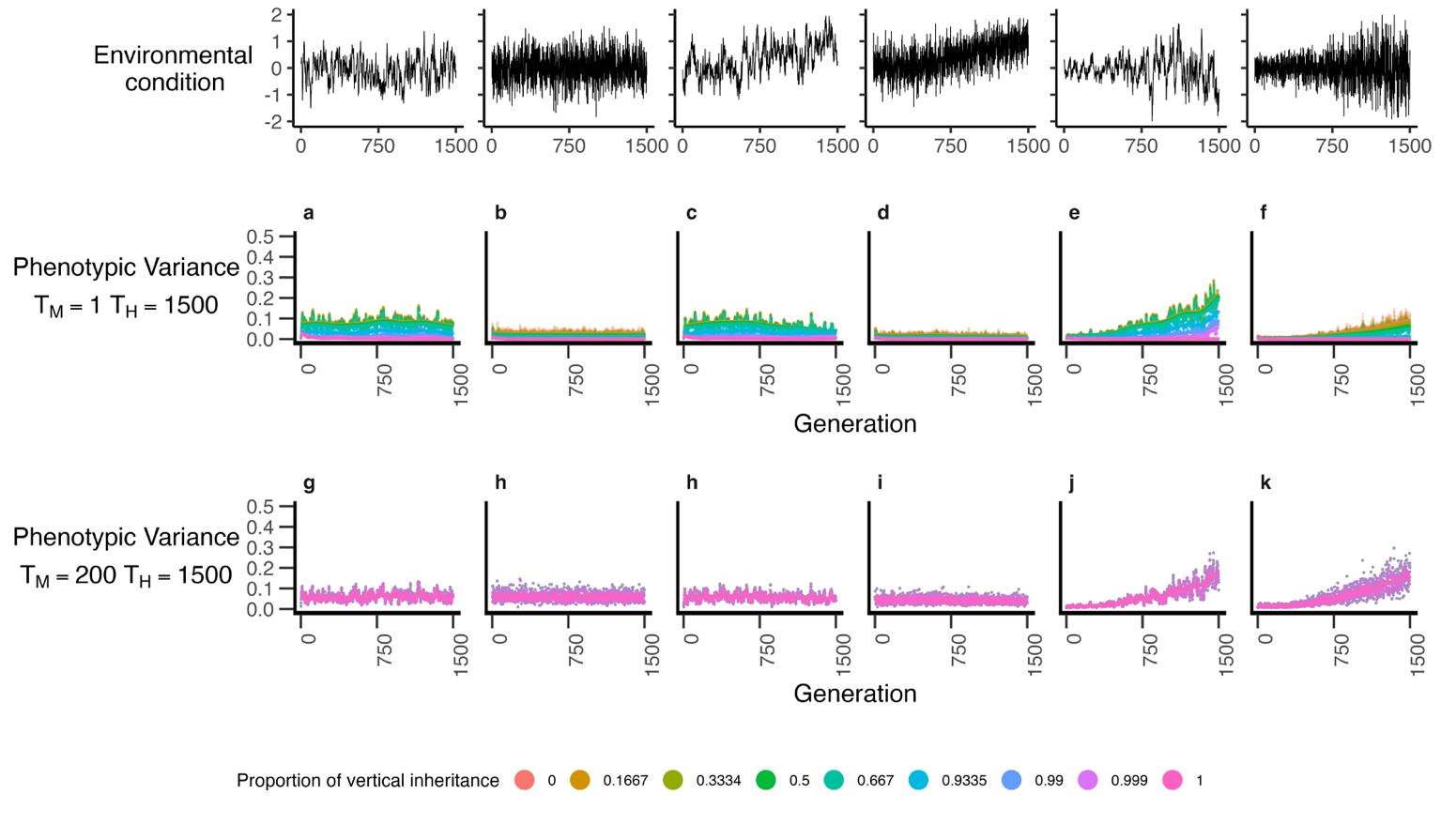
**Supplementary Material**

Supp. Fig. 1: Phenotypic variance of hosts over the course of our simulations where the number of host generations is 1(a-f) or is 200 (g-k). Lines represent GAM smoothing; data represents mean values from 20 replicate simulations.

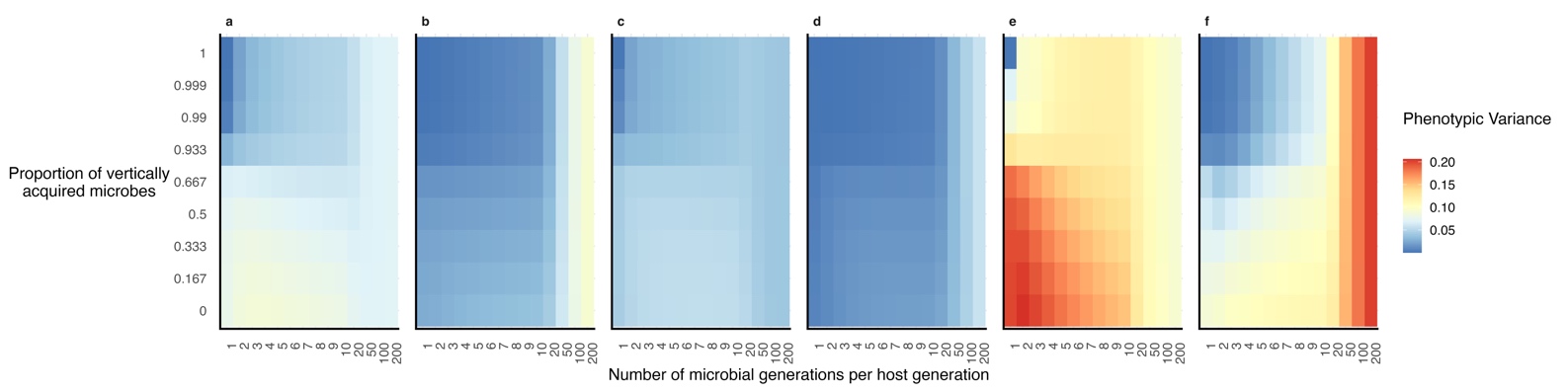

Supp. Fig. 2: Phenotypic variance of hosts at generation 1,500 in response to both vertical inheritance proportion and number of microbial generations per host generation (a-f).
